# Symbiotic *Escherichia coli* strains can better colonize host stinkbugs and outcompete natural symbiotic bacteria, but confer less fitness benefits

**DOI:** 10.64898/2026.04.01.715892

**Authors:** Wenjin Cai, Minoru Moriyama, Yudai Nishide, Ryuichi Koga, Takema Fukatsu

**Affiliations:** Molecular Biosystems Research Institute, National Institute of Advanced Industrial Science and Technology (AIST), Tsukuba, Japan; Department of Biological Sciences, Graduate School of Science, the University of Tokyo, Tokyo, Japan; Institute of Agrobiological Sciences Ohwashi, National Agriculture and Food Research Organization (NARO), Tsukuba, Japan; Graduate School of Life and Environmental Sciences, University of Tsukuba, Tsukuba, Japan

**Author notes:** Address correspondence to Minoru Moriyama, or Takema Fukatsu.

**Keywords:** *Plautia stali*, *Escherichia coli*, *Pantoea*, symbiont, competition, evolution, cheater

## Abstract

The stinkbug *Plautia stali* harbors essential gut symbiotic bacteria of the genus *Pantoea*, whose natural strains differ in cultivability and host benefits. Using this system, we evaluated how laboratory-evolved and genetically-engineered symbiotic *Escherichia coli* strains compete against native *Pantoea* symbionts and how they influence host fitness. In single infection assays, the native uncultivable symbiont Sym A conferred the highest host performance, whereas the evolved (CmL05G13) and artificial (Δ*cyaA*) symbiotic *E. coli* strains supported host survival at levels comparable to cultivable *Pantoea* symbionts (Sym C–F). In competitive co-infection assays, the symbiotic *E. coli* strains generally showed unexpectedly strong colonization ability. CmL05G13 outcompeted all the cultivable symbionts Sym C–F and even displaced the native uncultivable symbiont Sym A, whereas Δ*cyaA* and the nonsymbiotic control *E. coli* Δ*intS* were dominated by Sym A at the adult stage. Despite their superior infection competitiveness, the symbiotic *E. coli* strains provided limited reproductive benefits, behaving as “cheater-like” associates. They were able to invade and dominate the symbiotic organ but failed to match the fitness contributions of native symbionts. These results demonstrate that the experimentally evolved *E. coli* can rapidly acquire strong colonization ability surpassing that of the natural symbionts that have coevolved with *P. stali* in nature. At the same time, the mismatch between infection success and host fitness benefits highlights potential evolutionary conflicts and provides an experimental model for studying the dynamics of cheating, mutualism, and symbiont replacement in vertically transmitted symbioses.

**IMPORTANCE:** Understanding how novel symbionts invade and displace long-term mutualists is central to the evolution of symbiosis. This study demonstrates that *Escherichia coli*, originally a nonsymbiotic bacterium, can rapidly evolve potent colonization ability and even outcompete native *Pantoea* symbionts of the stinkbug *Plautia stali*. Meanwhile, these competitive *E. coli* strains confer markedly lower reproductive benefits compared with the native symbionts that have developed intimate mutualistic association with host *P. stali* over evolutionary time, revealing a striking decoupling between infection success and host fitness. This finding highlights the potential for cheater-like microbes to invade vertically transmitted symbioses and destabilize coevolved partnerships. By combining experimental evolution, controlled co-infections, and quantitative analyses, the *P. stali*–*E. coli* experimental symbiotic system provides a powerful model for studying the mechanisms and evolutionary dynamics of mutualism, cheating, and symbiont replacement.

## INTRODUCTION

Many organisms are associated with specific and beneficial microbial partners, which significantly contribute to growth and survival of the host organisms. In some cases, the host and the symbiont cannot survive without the intimate partnership, forming an almost inseparable biological entity and manifesting an ultimate form of symbiosis (1–3). At the beginning, however, such an essential microbial partner must have originated from a free-living environmental microbe. The origins and processes leading to such elaborate symbiotic systems we currently observe in nature comprise an important but challenging issue in evolutionary biology, to which experimental evolutionary approaches will provide valuable insights (4, 5).

Recently, the stinkbug *Plautia stali* emerged as an excellent model system that enables experimental and evolutionary approaches to microbial symbiosis. *P. stali* has a specialized symbiotic organ consisting of numerous sac-like crypts arranged in four rows along the posterior midgut, whose inner cavities harbor a specific bacterial symbiont of the genus *Pantoea* (6–8). The symbiont is essential for growth and survival of the host insect and vertically transmitted to next generation via maternal smearing of symbiont-containing secretion on the egg surface upon oviposition, which newborn nymphs lick and ingest (9, 10). Notably, the symbiotic bacteria of *P. stali* exhibit geographic polymorphism in Japan: an uncultivable symbiont *Pantoea* sp. A is fixed in mainland populations, whereas an uncultivable symbiont *Pantoea* sp. B predominates and coexists with cultivable symbionts *Pantoea* spp. C–F in southwestern island populations (9). The situation that both uncultivable and cultivable essential symbionts coexist in natural populations of the same insect species has enabled various experimental approaches to symbiosis (9, 11).

Furthermore, we developed an experimental symbiotic system consisting of *P. stali* as host and the model bacterium *Escherichia coli* as symbiont. We generated symbiont-eliminated newborn nymphs of *P. stali*, inoculated a hypermutating *E. coli* strain to them, and maintained them in the laboratory. Then, within several months to a year, some *E. coli*-infected evolutionary lines of *P. stali* exhibited significantly improved adult emergence rate and body color, showing rapid evolution of “mutualistic” *E. coli* strains in the laboratory (12). By making use of the *P. stali-E. coli* artificial symbiotic system, we have obtained laboratory-evolved symbiotic *E. coli* strains, and by identifying and engineering bacterial genes responsible for the symbiotic phenotypes, we have generated artificial symbiotic *E. coli* strains such as Δ*cyaA,* Δ*crp*, Δ*tnaA*, Δ*trpR*, Δ*metJ*, etc. (12–14).

Microbial competitions are expected to universally occur wherever multiple microbes coexist and exploit limited resources such as nutrients, space, ecological niches, etc. (15). Through single-and mixed-culture experiments in laboratory settings, a variety of interactions, processes and mechanisms underlying the formation of microbial communities have been investigated (16). For example, previous studies on other stinkbug symbiotic systems, in which their beneficial gut bacteria of the genus *Caballeronia* (or *Burkholderia sensu lato*) are environmentally acquired and cultivable, have uncovered some notable aspects of intra-host symbiont-symbiont interactions that shape the final symbiotic microbiota and specificity in the host insects. In *Riptortus pedestris*, many environmental *Caballeronia* strains can colonize and benefit hosts in single inoculation, but in co-inoculation, the native symbiont strain consistently outcompetes other strains, which accounts for the high symbiont specificity observed in natural populations of *R. pedestris* (17). In *R. pedestris* and *Coreus marginatus*, different *Caballeronia* strains representing different subclades can colonize the host insects alone, but they show strong competitive hierarchies in pairwise infections, which determine fine-scale host–symbiont specificities (18). In *Anasa tristis*, co-infection with multiple *Caballeronia* strains is commonly observed and dominant microbial strains show considerable diversity among different host individuals, in which the symbiont diversity/heterogeneity is promoted by bacterial population bottleneck upon nymphal acquisition and governed by ecological drift (19). Overall, these studies demonstrated that, for each symbiotic system, the native symbiont strain tends to, if not always, outcompetes other strains and establishes the specific symbiotic association, wherein symbiont-symbiont competition as well as symbiont population bottleneck play important roles. Here a question emerges – How competitive are the evolved and artificial symbiotic *E. coli* strains against the native symbiont strains of *P. stali* when co-inoculated to the host insects? Intuitively, it is expected that, even if the symbiotic *E. coli* strains have become mutualistic to *P. stali*, the laboratory evolution of the model bacterium within a year or so cannot reach the level of symbiotic capability that has been attained by the native symbiont strains over evolutionary time under natural conditions. However, we found that, unexpectedly, the symbiotic *E. coli* strains tend to outcompete the native symbiotic bacteria when co-inoculated to *P. stali*. On the other hand, despite the superior infection competitiveness, fecundity levels of the host insects infected with the symbiotic *E. coli* strains were significantly lower than those with the native symbiont strains. These findings highlight that the symbiotic *E. coli* strains contribute to different host fitness components in different ways, thereby potentially behaving as cheater-like microbial associates for *P. stali*.

## RESULTS

### Experimental insects, symbiotic bacteria, *E. coli* strains, and their phenotypes

In this study, all the experiments were conducted using host insects of the same genetic background, an inbred strain of *P. stali* that was originally collected at Tsukuba, Ibaraki, Japan. The native symbiotic bacterium of the insect strain was *Pantoea* sp. A (hereafter Sym A), which is yellow in color and uncultivable (Fig. 1A-C). The following bacterial strains were inoculated to aseptic newborn nymphs of *P. stali* that were generated by sterilization of egg surface (20). The cultivable symbiotic bacteria, *Pantoea* spp. C, D, E and F (hereafter Sym C, Sym D, Sym E and Sym F), were originally isolated from southwestern island populations of *P. stali* (9), which are all cultivable, form yellow colonies on agar plates, and confer yellow coloration to the symbiotic organ of their hosts (Fig. 1D-O). The control nonsymbiotic *E. coli* strain Δ*intS*, the evolved symbiotic *E. coli* strain CmL05G13 and the artificial symbiotic *E. coli* strain Δ*cyaA* were obtained and used in previous experimental evolutionary studies (12–14, 21), which are cultivable, form white colonies on agar plates. confer white coloration to the symbiotic organ of their host insects (Fig. 1P-X). Note that colonies of the symbiotic *E. coli* strains CmL05G13 and Δ*cyaA* are smaller due to attenuated growth rates, which are probably ascribed to evolutionary trade-off (Fig. 1U, X) (12).

**FIG 1.**
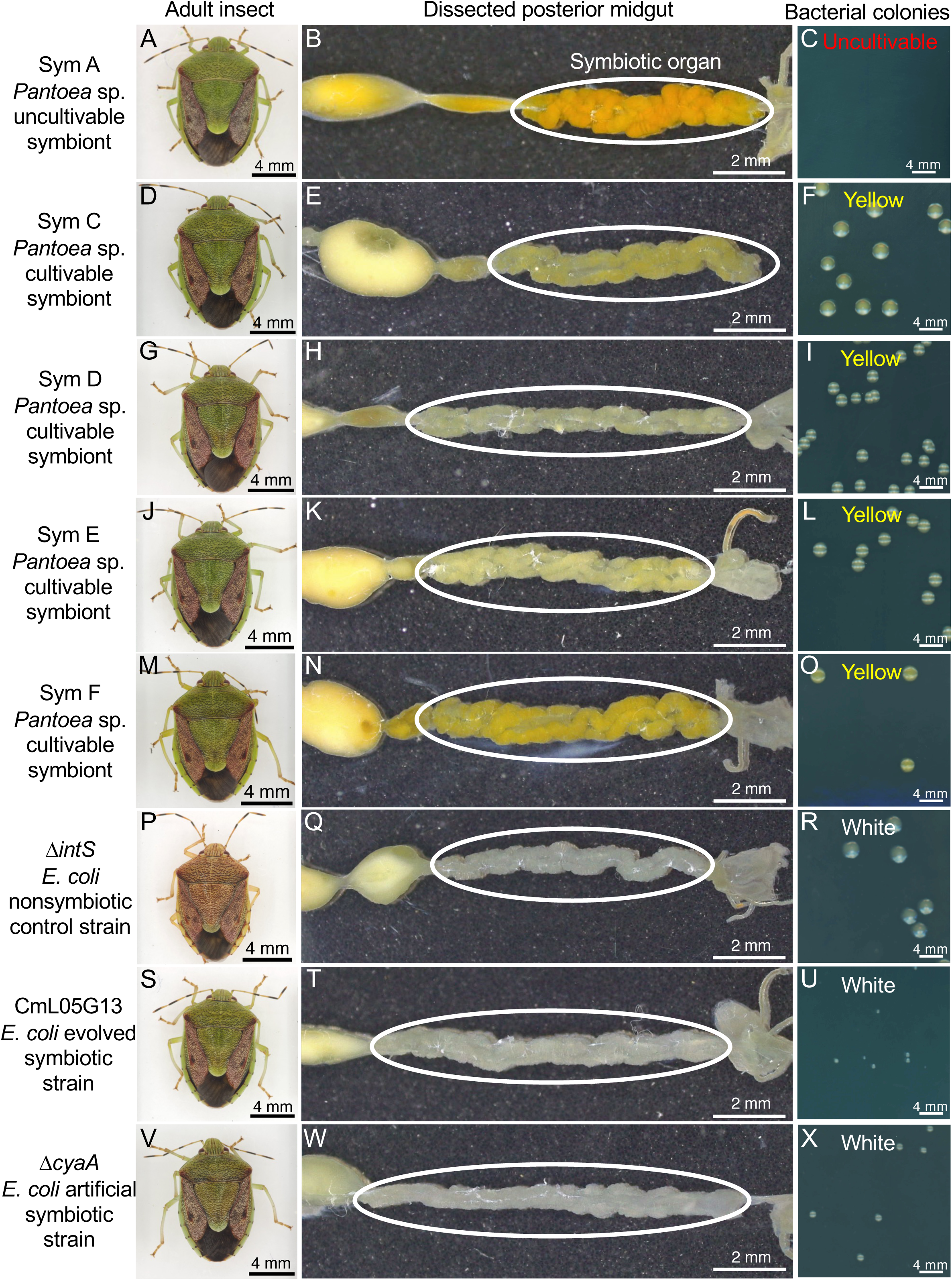
Host insects, symbiotic organs and bacteria examined in this study. (**A-C**) *P. stali* originally infected with the natural uncultivable symbiont *Pantoea* sp. A (Sym A). (**D-F**) *P. stali* experimentally infected with the natural cultivable symbiont *Pantoea* sp. C (Sym C). (**G-I**) *P. stali* experimentally infected with the natural cultivable symbiont *Pantoea* sp. D (Sym D). (**J-L**) *P. stali* experimentally infected with the natural uncultivable symbiont *Pantoea* sp. E (Sym E). (**M-O**) *P. stali* experimentally infected with the natural uncultivable symbiont *Pantoea* sp. F (Sym F). (**P-R**) *P. stali* experimentally infected with the control nonsymbiotic *E. coli* strain Δ*intS*. (**S-U**) *P. stali* experimentally infected with the evolved symbiotic *E. coli* strain CmL05G13. (**V-X**) *P. stali* experimentally infected with the artificial symbiotic *E. coli* strain Δ*cyaA*. (**A, D, G, J, M, P, S, V**) Adult insects. Note that Δ*intS*-infected insect is darker in color and smaller in size. (**B, E, H, K, N, Q, T, W**) Dissected posterior midguts. The symbiotic organ is circled, whose color is yellow for *Pantoea* infections and white for *E. coli* infections, respectively. (**C, F, I, L O, R, U, X**) Bacterial colonies cultured on LB agar plates at 25°C for 48 h. Note that colony color is yellow for cultivable *Pantoea* symbionts and white for *E. coli* strains.

### Adult emergence rate of single-infected insects with uncultivable/cultivable symbiotic bacteria of *P. stali* and nonsymbiotic/symbiotic *E. coli* strains

The insects harboring the native uncultivable symbiont Sym A attained the highest adult emergence rate of 71.7% on average. Among the insects infected with the cultivable symbionts, while the insects infected with Sym C showed a low adult emergence rate of 19.5% on average, the insects infected with Sym D, Sym E and Sym F exhibited remarkably high adult emergence rates of 46.1%, 55.8% and 56.1%, respectively. The insects infected with the control nonsymbiotic *E. coli* Δ*intS* showed the lowest adult emergence rate of 3.9% on average, whereas the insects infected with the evolved symbiotic *E. coli* CmL05G13 and the artificial symbiotic *E. coli* Δ*cyaA* exhibited remarkably high adult emergence rates of 56.5% and 56.0%, respectively. Hence, the order of the adult emergence rates was Sym A > Sym D, Sym E, Sym F, CmL05G13, Δ*cyaA* > Sym C > Δ*intS* (Fig. 2A).

**FIG 2.**
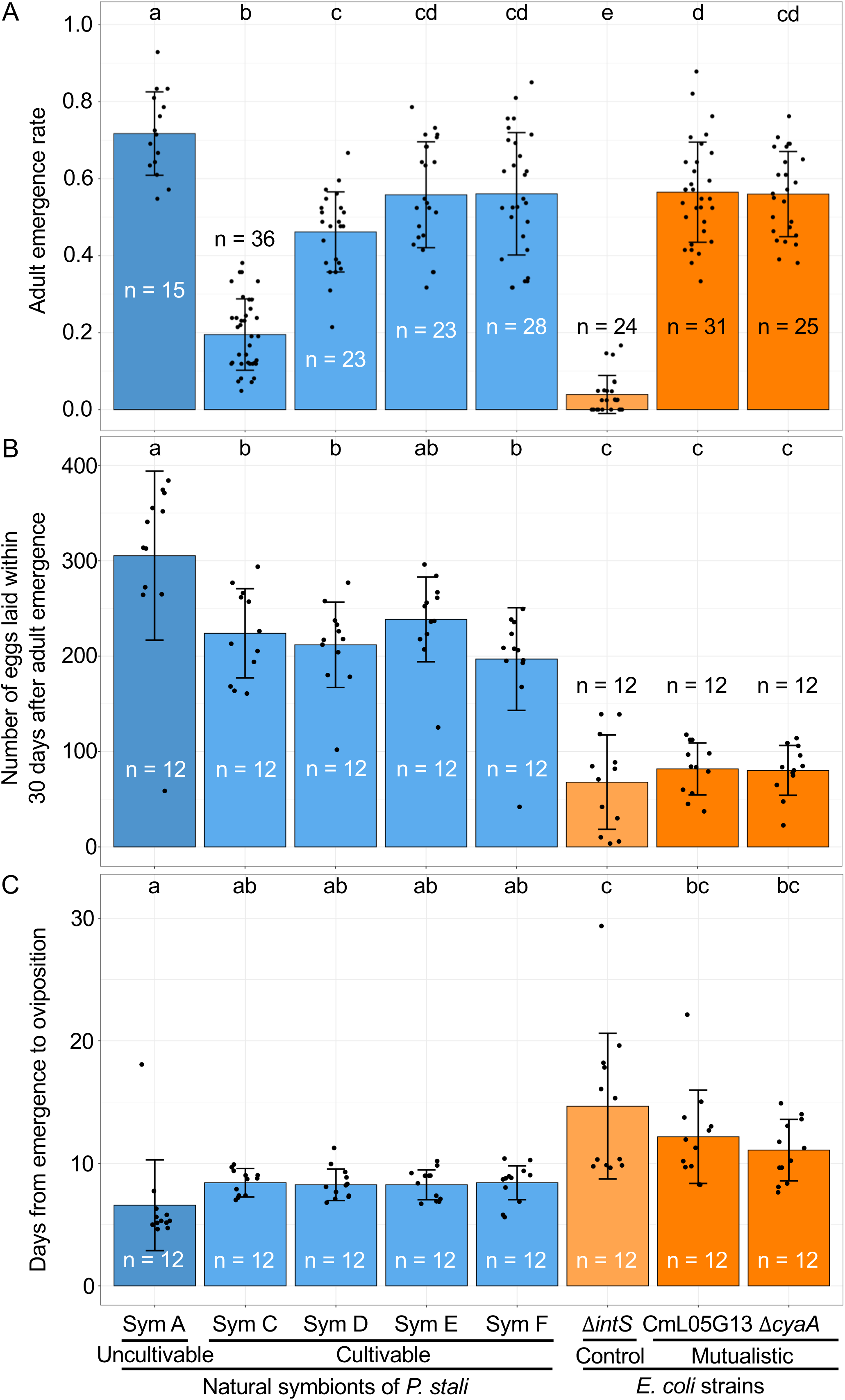
Fitness effects of single-infections with uncultivable/cultivable symbiotic bacteria and nonsymbiotic/symbiotic *E. coli* strains on *P. stali*. (**A**) Adult emergence rate. (**B**) Number of eggs laid within 30 days after adult emergence. (**C**) Days from adult emergence to first oviposition. Bars, whiskers and dots indicate means, standard deviations, and data points, respectively. Different alphabetical letters (a, b, c, d) indicate statistically significant differences (pairwise Wilcoxon rank-sum test with Hommel’s correction; *P* < 0.05).

### Fecundity of single-infected insects with uncultivable/cultivable symbiotic bacteria and nonsymbiotic/symbiotic *E. coli* strains

As for number of eggs laid within 30 days after adult emergence, the adult insects harboring the native uncultivable symbiont Sym A produced around 300 eggs on average, which was significantly more than some 200 eggs produced by the adult insects infected with the cultivable symbionts Sym C, Sym D, Sym E and Sym F. The adult insects infected with the nonsymbiotic and symbiotic *E. coli* strains Δ*intS,* CmL05G13 and Δ*cyaA* exhibited significantly lower host fecundity, yielding less than 100 eggs on average.

Hence, the order of the levels of host fecundity was Sym A > Sym C, Sym D, Sym E, Sym F > Δ*intS*, CmL05G13, Δ*cyaA* (Fig. 2B).

### Time to reproduction in single-infected insects with uncultivable/cultivable symbiotic bacteria and nonsymbiotic/symbiotic *E. coli* strains

As for days from adult emergence to first oviposition, the adult insects harboring the native uncultivable symbiont Sym A took mostly 5-6 days, whereas the adult insects infected with the cultivable symbionts Sym C, Sym D, Sym E and Sym F took around 8-10 days. The adult insects infected with the nonsymbiotic and symbiotic *E. coli* strains Δ*intS,* CmL05G13 and Δ*cyaA* took over 10 days, of which Δ*intS* exhibited the most remarkable delay. Hence, the order of the durations needed for host reproductive maturity was Sym A < Sym C, Sym D, Sym E, Sym F < CmL05G13, Δ*cyaA* < Δ*intS* (Fig. 2C).

### Evolved and artificial symbiotic *E. coli* strains are comparable to culturable symbiotic bacteria in supporting host growth and survival, but inferior in supporting host reproduction

Taken together, these results indicated that (i) the native uncultivable symbiont Sym A is the best symbiont as expected, attaining the highest adult emergence rate, the fastest reproductive maturity, and the highest fecundity, (ii) the control nonsymbiotic *E. coli* strain Δ*intS* exhibits the worst symbiotic performance as expected, suffering the lowest adult emergence rate, the latest reproductive maturity, and the lowest fecundity, and (iii) the evolved symbiotic *E. coli* strain CmL05G13 and the artificial symbiotic *E. coli* strain Δ*cyaA* can support the host growth pretty well, attaining adult emergence rates and reproductive maturity comparable to those with the cultivable symbiotic bacteria Sym C, Sym D, Sym E and Sym F, though not comparable to Sym A, but (iv) the symbiotic *E. coli* strains CmL05G13 and Δ*cyaA* show significantly lower host fecundity in comparison with the uncultivable and cultivable symbionts Sym A, Sym C, Sym D, Sym E and Sym F. Overall, it turned out that the symbiotic *E. coli* strains CmL05G13 and Δ*cyaA* are certainly capable of improving host growth and survival, but improvement of host fecundity is marginal.

### Bacterial titers in single-infected insects with uncultivable/cultivable symbiotic bacteria and nonsymbiotic/symbiotic *E. coli* strains

In second instar nymphs, the native uncultivable symbiont Sym A, the evolved symbiotic *E. coli* CmL05G13, and the artificial symbiotic *E. coli* Δ*cyaA* exhibited relatively high bacterial titers around 10^6^ gene copies equivalent, whereas the cultivable symbionts Sym C, Sym D, Sym E and Sym F, and the control nonsymbiotic *E. coli* Δ*intS* showed relatively low bacterial titers around 10^4^ gene copies equivalent (Fig. 3A). In adult insects, similar patterns were observed. The native uncultivable symbiont Sym A, the evolved symbiotic *E. coli* CmL05G13, and the artificial symbiotic *E. coli* Δ*cyaA* exhibited relatively high bacterial titers around 10^8^ gene copies equivalent, whereas the cultivable symbionts Sym C, Sym D, Sym E and Sym F, and the control nonsymbiotic *E. coli* Δ*intS* showed relatively low bacterial titers around 10^6^ gene copies equivalent (Fig. 3B). These patterns revealed that (i) among the symbiotic bacteria, the levels of bacterial titers are positively correlated with the levels of beneficial effects on the host insects, (ii) among the *E. coli* strains, the levels of bacterial titers are also positively correlated with the levels of beneficial effects on the host insects, and thus (iii) the elevated bacterial titers may underpin the improved performance of the infected host insects.

**FIG 3.**
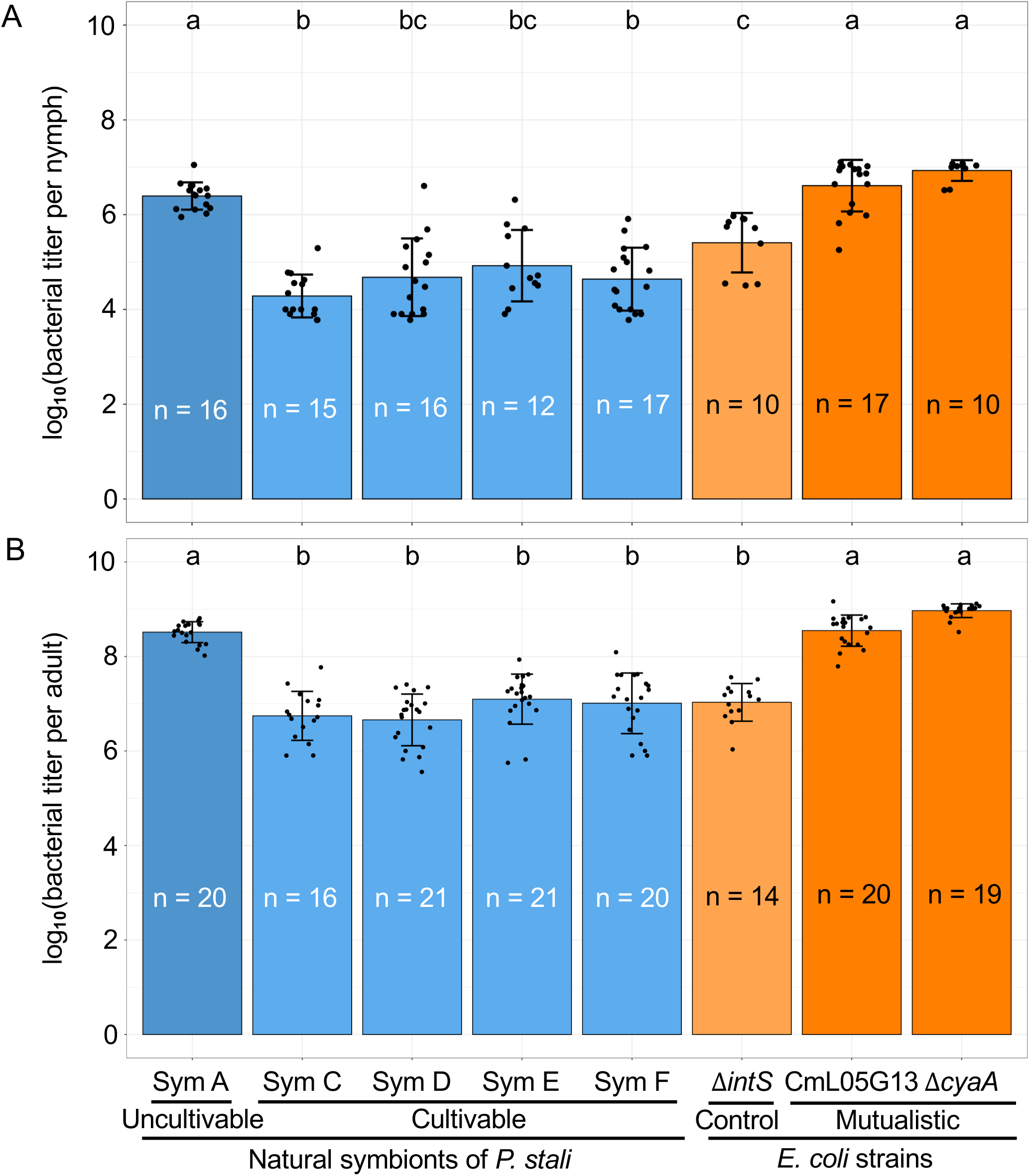
Bacterial titers of single-infected uncultivable/cultivable symbiotic bacteria and nonsymbiotic/symbiotic *E. coli* strains in *P. stali*. (**A**) Bacterial titer in second instar nymph. (**B**) Bacterial titer in adult. Note that the vertical axes are log-scaled. Bars, whiskers and dots indicate means, standard deviations and data points, respectively. Different alphabetical letters (a, b, c) indicate statistically significant differences (pairwise Wilcoxon rank-sum test with Hommel’s correction; *P* < 0.05).

### Competitive outcomes of co-infection with cultivable symbiotic bacterium Sym C and nonsymbiotic/symbiotic *E. coli* strains

We performed competitive infection experiments between the cultivable symbiont Sym C and the nonsymbiotic/symbiotic *E. coli* strains by inoculating 1:1 mixture of Sym C cells and *E. coli* cells to aseptic newborn nymphs of *P. stali*. In this series of experiments, bacterial detection and quantification were conducted by CFU-based methods.

#### Sym C vs nonsymbiotic control *E. coli* Δ*intS*

When Sym C cells and Δ*intS* cells were co-inoculated, Sym C preferentially established infection in the host symbiotic organ over Δ*intS* (Fig. 4; Fig. 5A). The adult emergence rates of the co-inoculated insects were comparable to those of Sym C-infected insects and significantly higher than those of Δ*intS*-infected insects (Fig. 5B), which probably reflect the predominant establishment of Sym C in the co-inoculated adult insects (Fig. 4B). On the other hand, the bacterial titers in the co-inoculated insects were comparable to those in the Sym C-infected insects as well as those in the Δ*intS*-infected insects at the adult stage, although they significantly differed at the second instar stage (Fig. 5C, D). These results indicated that the cultivable symbiont Sym C is superior to the nonsymbiotic *E coli* Δ*intS* in both infection competitiveness and host survival improvement.

**FIG 4.**
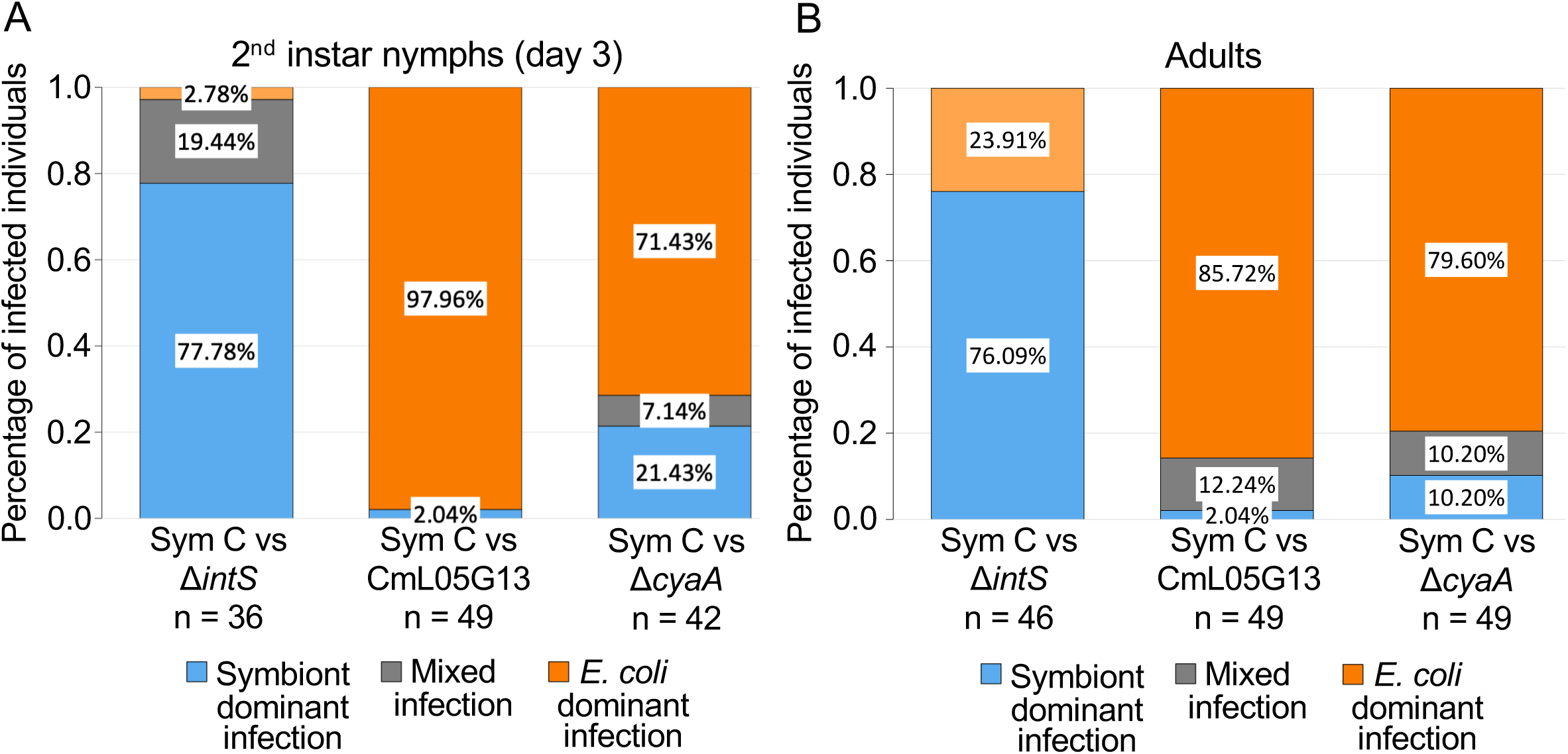
Bacterial predominance in *P. stali* co-inoculated with cultivable symbiotic bacterium Sym C and nonsymbiotic/symbiotic *E. coli* strains. (**A**) Second instar nymphs three days after molting. Each whole nymph was homogenized, diluted and plated, and subjected to observation of bacterial colonies. (**B**) Adult insects. Each dissected midgut symbiotic organ was homogenized, diluted and plated, and subjected to observation of bacterial colonies. Each plate with about 30-500 colonies was categorized into either “symbiont dominant infection” in which all colonies are yellow Sym C colonies (blue), “*E. coli* dominant infection” in which all colonies are white *E. coli* colonies (orange), and “mixed infection” in which both yellow Sym C colonies and white *E. coli* colonies are observed (gray).

**FIG 5.**
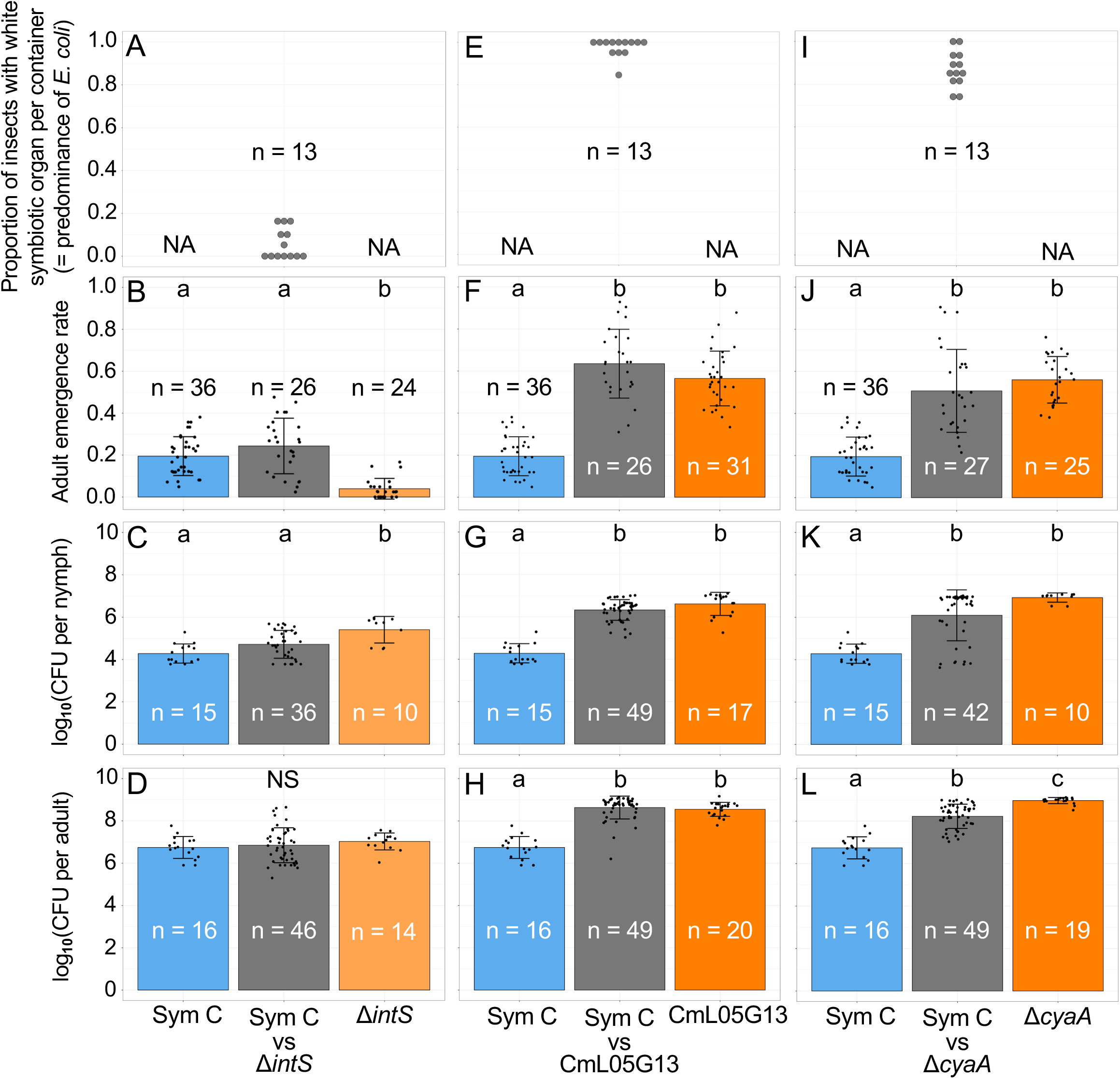
Bacterial predominance, adult emergence rate, and bacterial titer in *P. stali* co-inoculated with cultivable symbiotic bacterium Sym C and nonsymbiotic/symbiotic *E. coli* strains. (**A-D**) Co-inoculation with Sym C and the nonsymbiotic control *E. coli* Δ*intS*. (**E-H**) Co-inoculation with Sym C and the evolved symbiotic *E. coli* CmL05G13. (**I-L**) Co-inoculation with Sym C and the artificial symbiotic *E. coli* Δ*cyaA*. (**A, E, I**) Proportion of adult insects with white symbiotic organ per rearing container, each containing 3-32 insects, indicating the level of *E. coli* predominance in the co-inoculated host insects. (**B, F, J**) Adult emergence rates of Sym C-inoculated, co-inoculated and *E. coli*-inoculated insects. (**C, G, K**) Bacterial titers in Sym C-inoculated, co-inoculated and *E. coli*-inoculated second instar nymphs in terms of CFU per nymph. (**D, H, L**) Bacterial titers in Sym C-inoculated, co-inoculated and *E. coli*-inoculated adult insects in terms of CFU per dissected symbiotic organ. In (**B-D**), (**F-H**) and (**J-L**), blue, gray, and orange indicate Sym C-inoculated (with adult emergence rates and bacterial titers shown in Fig. 2 and Fig. 3), co-inoculated, and *E. coli*-inoculated (with adult emergence rates and bacterial titers shown in Fig. 2 and Fig. 3), respectively. Bars, whiskers and dots indicate means, standard deviations and data points, respectively. Different alphabetical letters (a, b, c) indicate statistically significant differences (pairwise Wilcoxon rank-sum test with Hommel’s correction; *P* < 0.05).

#### Sym C vs evolved symbiotic *E. coli* CmL05G13

When Sym C cells and CmL05G13 cells were co-inoculated, CmL05G13 predominantly established infection in the host symbiotic organ over Sym C (Fig. 4; Fig. 5E). The adult emergence rates of the co-inoculated insects were comparable to those of CmL05G13-infected insects and significantly higher than those of Sym C-infected insects (Fig. 5F), which probably reflect the preferential establishment of CmL05G13 in the co-inoculated adult insects (Fig. 4B). The bacterial titers in the co-inoculated insects were comparable to those in the CmL05G13-infected insects and significantly higher than those in the Sym C-infected insects (Fig. 5G, H). These results indicated that the evolved symbiotic *E. coli* CmL05G13 is superior to the cultivable symbiont Sym C in both infection competitiveness and host survival improvement.

#### Sym C vs artificial symbiotic *E. coli* Δ*cyaA*

When Sym C cells and Δ*cyaA* cells were co-inoculated, Δ*cyaA* predominantly established infection in the host symbiotic organ over Sym C (Fig. 4; Fig. 5I). The adult emergence rates of the co-inoculated insects were comparable to those of Δ*cyaA*-infected insects and significantly higher than those of Sym C-infected insects (Fig. 5J), which probably reflect the preferential establishment of Δ*cyaA* in the co-inoculated adult insects (Fig. 4B). The bacterial titers in the co-inoculated insects were comparable to those in the Δ*cyaA*-infected insects and significantly higher than those in the Sym C-infected insects (Fig. 5K, L). These results indicated that the artificial symbiotic *E. coli* Δ*cyaA* is superior to the cultivable symbiont Sym C in both infection competitiveness and host survival improvement.

Taken together, the cultivable symbiont Sym C isolated from a natural population of *P. stali* certainly shows better symbiotic capability than the control nonsymbiotic *E. coli* Δ*intS*, but, strikingly, the symbiotic *E. coli* CmL05G13 obtained through experimental symbiotic evolution and the artificial symbiotic *E. coli* generated by single-gene knockout exhibit better symbiotic capability than Sym C.

#### Competitive outcomes of co-infection with cultivable symbiotic bacteria Sym D/Sym E/Sym F and evolved symbiotic *E. coli* strain CmL05G13

Next, we performed competitive infection experiments between the cultivable symbiont Sym D/Sym E/Sym F and the evolved symbiotic *E. coli* strain CmL05G13 by inoculating 1:1 mixture of symbiont cells and *E. coli* cells to aseptic newborn nymphs of *P. stali*. In this series of experiments, bacterial detection and quantification were conducted by CFU-based methods.

#### Sym D vs evolved symbiotic *E. coli* CmL05G13

When Sym D cells and CmL05G13 cells were co-inoculated, CmL05G13 outcompeted Sym D almost completely (Fig. 6; Fig. 7A). The adult emergence rates of the co-inoculated insects were statistically neither different from those of the CmL05G13-infected insects nor those of the Sym D-infected insects, whereas the adult emergence rates of the CmL05G13-infected insects were significantly higher than those of the Sym D-infected insects (Fig. 7B). On the other hand, the bacterial titers in the co-inoculated insects were, though comparable to or somewhat lower than those in the CmL05G13-infected insects, significantly higher than those in the Sym D-infected insects (Fig. 7C, D). These results indicated that the evolved symbiotic *E. coli* CmL05G13 is superior to the cultivable symbiont Sym D in both infection competitiveness and host survival improvement.

**FIG 6.**
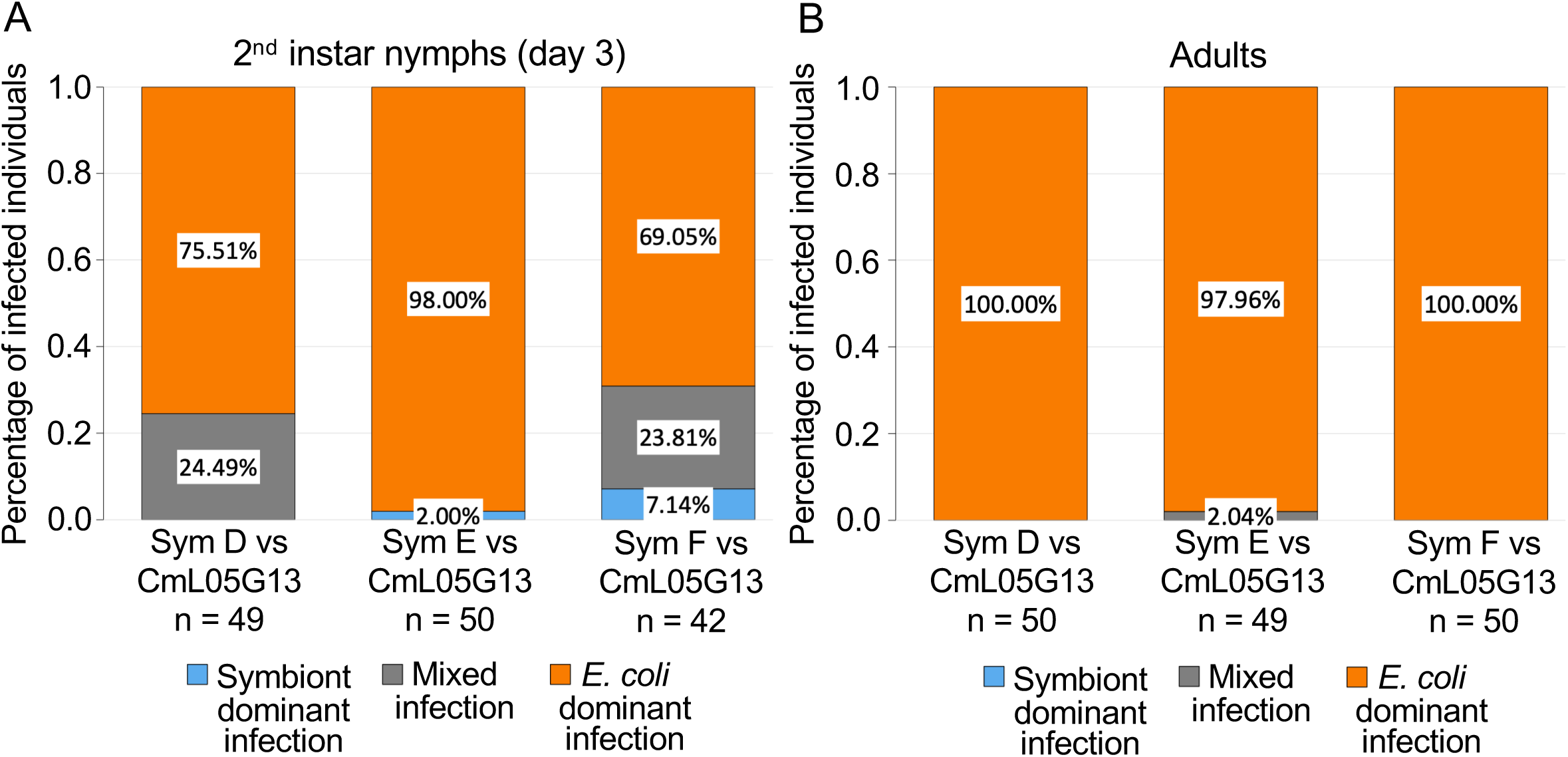
Bacterial predominance in *P. stali* co-inoculated with cultivable symbiotic bacteria Sym D/Sym E/Sym F and evolved symbiotic *E. coli* strain CmL05G13. (**A**) Second instar nymphs three days after molting. Each whole nymph was homogenized, diluted and plated, and subjected to observation of bacterial colonies. (**B**) Adult insects. Each dissected midgut symbiotic organ was homogenized, diluted and plated, and subjected to observation of bacterial colonies. Each plate with about 30-500 colonies was categorized into either “symbiont dominant infection” in which all colonies are yellow Sym C colonies (blue), “*E. coli* dominant infection” in which all colonies are white *E. coli* colonies (orange), and “mixed infection” in which both yellow Sym C colonies and white *E. coli* colonies are observed (gray).

**FIG 7.**
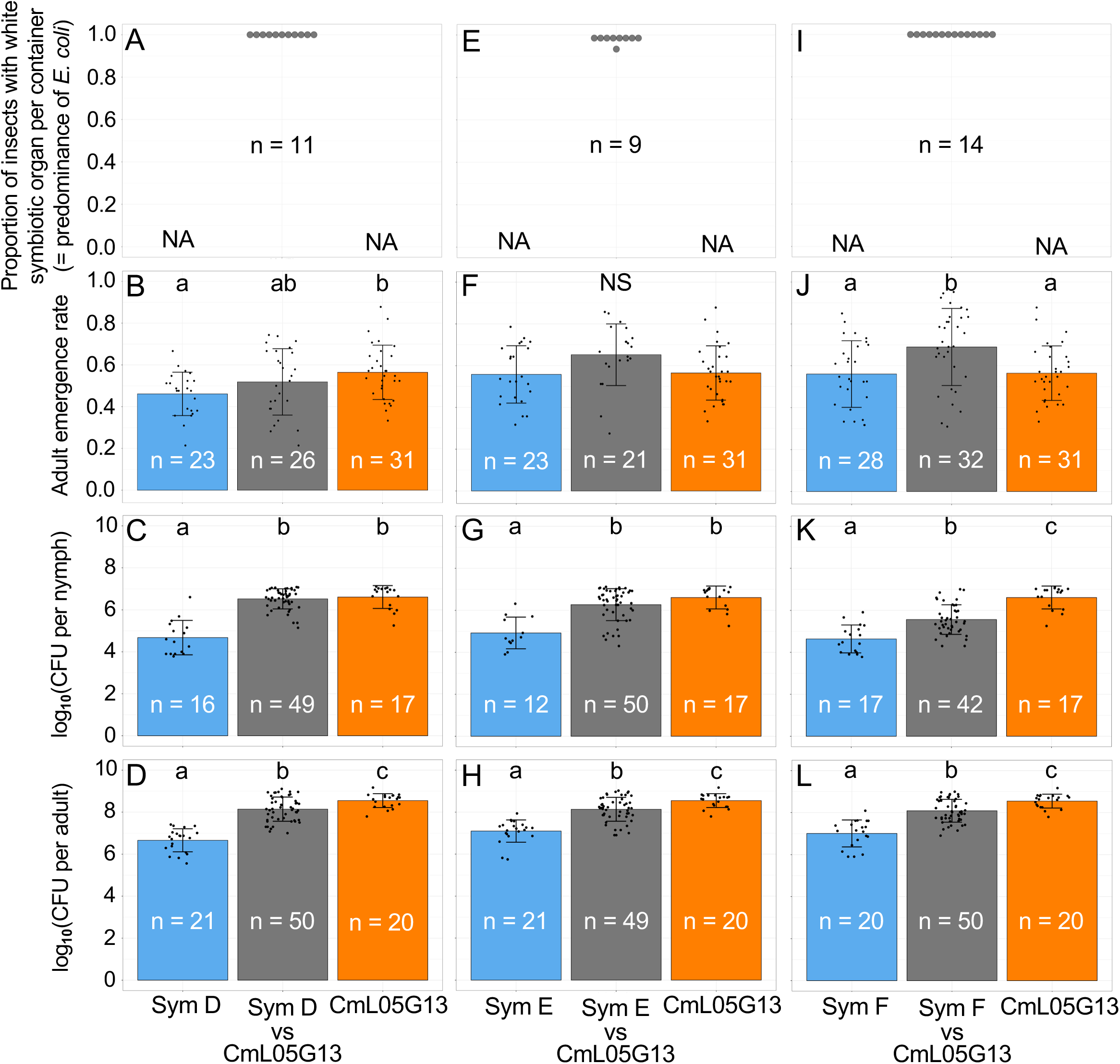
Bacterial predominance, adult emergence rate, and bacterial titer in *P. stali* co-inoculated with cultivable symbiotic bacteria Sym D/Sym E/SymF and evolved symbiotic *E. coli* strain CmL05G13. (**A-D**) Co-inoculation with Sym D and CmL05G13. (**E-H**) Co-inoculation with Sym E and CmL05G13. (**I-L**) Co-inoculation with Sym F and CmL05G13. (**A, E, I**) Proportion of adult insects with white symbiotic organ per rearing container, each containing 9-40 insects, indicating the level of *E. coli* predominance in the co-inoculated host insects. (**B, F, J**) Adult emergence rates of Sym D/Sym E/Sym F-inoculated, co-inoculated and CmL05G13-inoculated insects. (**C, G, K**) Bacterial titers in Sym D/Sym E/SymF-inoculated, co-inoculated and CmL05G13-inoculated second instar nymphs in terms of CFU per nymph. (**D, H, L**) Bacterial titers in Sym D/Sym E/Sym F-inoculated, co-inoculated and CmL05G13-inoculated adult insects in terms of CFU per dissected symbiotic organ. In (**B-D**), (**F-H**) and (**J-L**), blue, gray, and orange indicate Sym D/Sym E/Sym F-inoculated (with adult emergence rates and bacterial titers shown in Fig. 2 and Fig. 3), co-inoculated, and CmL05G13-inoculated (with adult emergence rates and bacterial titers shown in Fig. 2 and Fig. 3), respectively. Bars, whiskers and dots indicate means, standard deviations and data points, respectively. Different alphabetical letters (a, b, c) indicate statistically significant differences (pairwise Wilcoxon rank-sum test with Hommel’s correction; *P* < 0.05).

#### Sym E vs evolved symbiotic *E. coli* CmL05G13

When Sym E cells and CmL05G13 cells were co-inoculated, CmL05G13 outcompeted Sym E almost completely (Fig. 6; Fig. 7E). The adult emergence rates of the co-inoculated insects were statistically neither different from those of CmL05G13-infected insects nor those of Sym E-infected insects (Fig. 7F). The bacterial titers in the co-inoculated insects were, while comparable to or somewhat lower than the CmL05G13-infected insects, significantly higher than those in the Sym E-infected insects (Fig. 7G, H). These results indicated that the evolved symbiotic *E. coli* CmL05G13 is superior to the cultivable symbiont Sym E in infection competitiveness.

#### Sym F vs evolved symbiotic *E. coli* CmL05G13

When Sym F cells and CmL05G13 cells were co-inoculated, CmL05G13 outcompeted Sym F almost completely (Fig. 6; Fig. 7I). The adult emergence rates of the co-inoculated insects were significantly higher than those of the CmL05G13-infected insects and those of the Sym F-infected insects (Fig. 7J). The bacterial titers in the co-inoculated insects were, while lower than those in the CmL05G13-infected insects, higher than those in the Sym F-infected insects (Fig. 7K, L). These results indicated that the evolved symbiotic *E. coli* CmL05G13 is superior to the cultivable symbiont Sym F in infection competitiveness.

Taken together, the symbiotic *E. coli* CmL05G13, which was obtained through experimental symbiotic evolution with *P. stali* in the laboratory, shows better symbiotic capability than the cultivable symbiotic bacteria Sym D, Sym E and Sym F that were isolated from natural populations of *P. stali*.

### Competitive outcomes of co-infection with native uncultivable symbiotic bacterium Sym A and nonsymbiotic/symbiotic *E. coli* strains

Finally, we performed competitive infection experiments between the native uncultivable symbiont Sym A and the nonsymbiotic/symbiotic *E. coli* strains by inoculating approximately 1:1 mixture of Sym A cells and *E. coli* cells to aseptic newborn nymphs of *P. stali*. In this series of experiments, bacterial detection and quantification were conducted by quantitative PCR because Sym A is uncultivable.

#### SymA vs nonsymbiotic control *E. coli* Δ*intS*

When Sym A cells and Δ*intS* cells were co-inoculated, Δ*intS* established predominant infection over Sym A at the second instar stage (Fig. 8A). However, the infection predominance was reversed at the adult stage, when Sym A almost outcompeted Δ*intS* (Fig. 8B; Fig. 9A). The adult emergence rates of the co-inoculated insects were lower but close to those of Sym A-infected insects, and were drastically higher than those of Δ*intS*-infected insects (Fig. 9B), which must reflect the predominant establishment of Sym A in the co-inoculated adult insects (Fig. 8B). The bacterial titers in the co-inoculated insects were lower or comparable to those in the Sym A-infected insects and significantly higher than those in the Δ*intS*-infected insects (Fig. 9C, D). These results indicated that, although the nonsymbiotic *E. coli* Δ*intS* may be good at initial infection and proliferation, the native uncultivable symbiont Sym A is superior in establishing the symbiosis with its original host *P. stali* and finally excludes the competitor *E. coli*, thereby attaining improved host survival.

**FIG 8.**
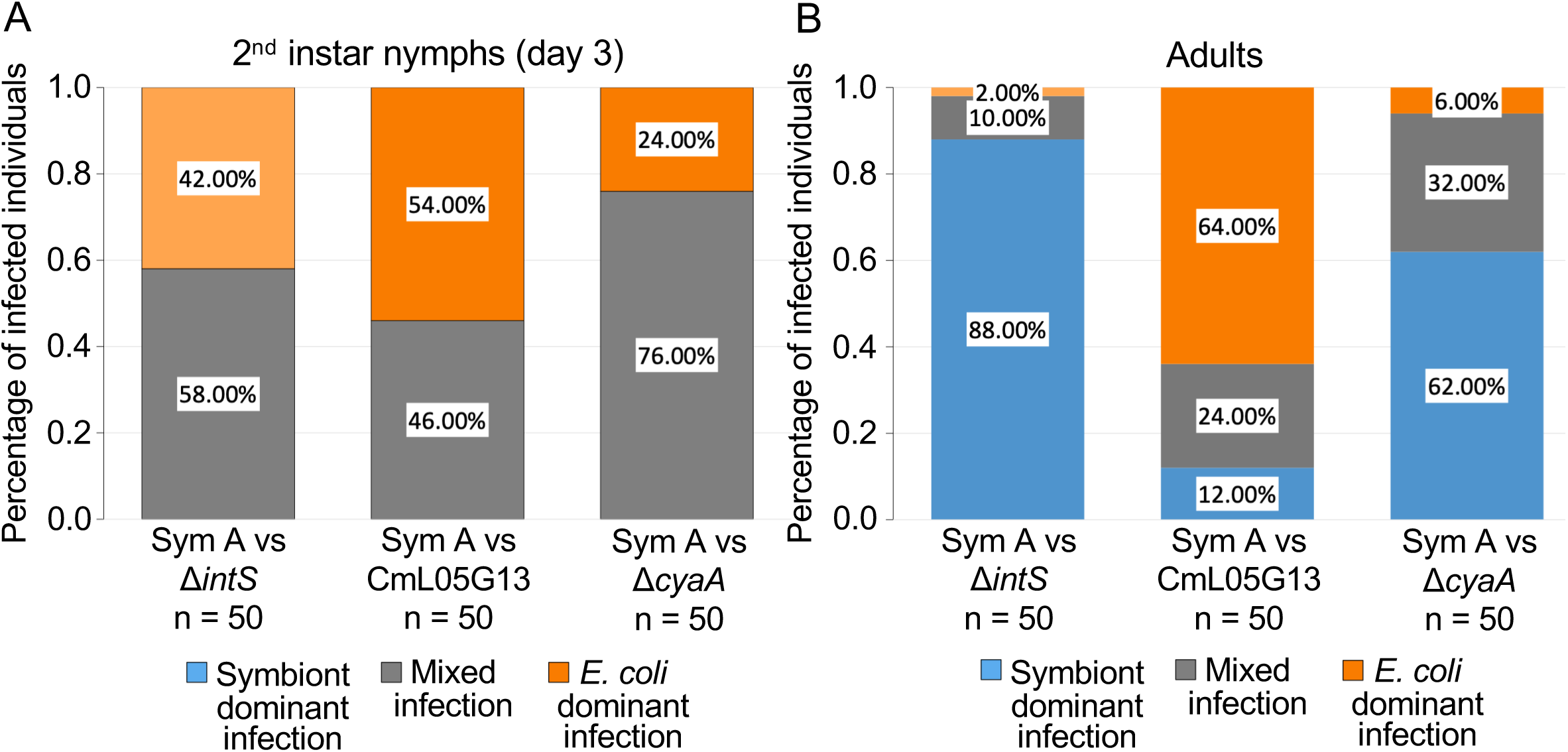
Bacterial predominance in *P. stali* co-inoculated with uncultivable symbiotic bacterium Sym A and nonsymbiotic/symbiotic *E. coli* strains. (**A**) Second instar nymphs three days after molting. Each whole nymph was subjected to DNA extraction and quantitative PCR targeting *groEL* gene of Sym A and *E. coli*. (**B**) Adult insects. Each dissected midgut symbiotic organ was subjected to DNA extraction and quantitative PCR of *groEL* gene of Sym A and *E. coli*. Each insect was categorized into either “symbiont dominant infection” in which Sym A titer is more than 95% of total bacterial titer (blue), “*E. coli* dominant infection” in which *E. coli* titer is more than 95% of total bacterial titer (orange), and “mixed infection” in which neither Sym A nor *E coli* is more than 95% of total bacterial titer (gray).

**FIG 9.**
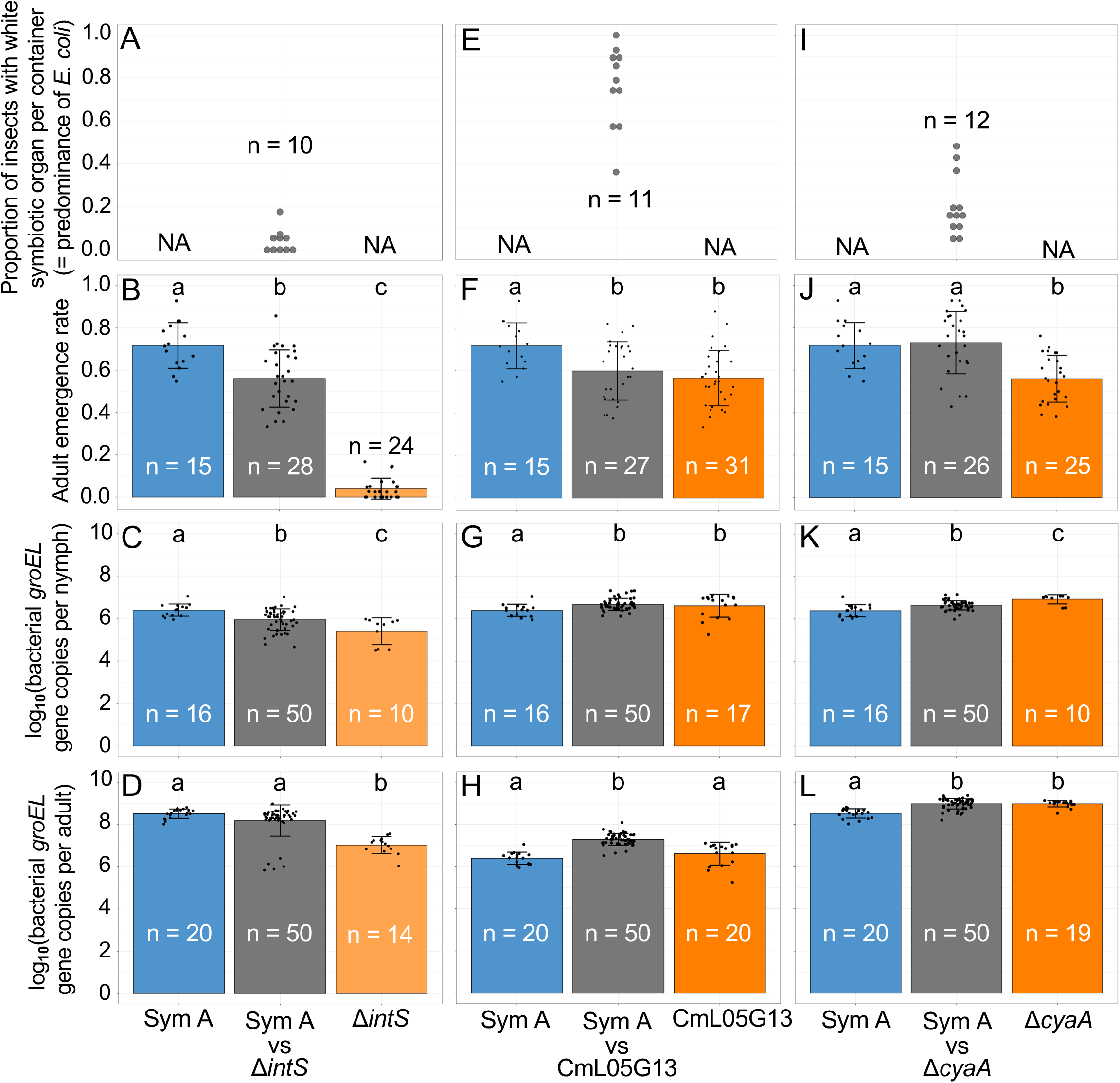
Bacterial predominance, adult emergence rate, and bacterial titer in *P. stali* co-inoculated with uncultivable symbiotic bacterium Sym A and nonsymbiotic/symbiotic *E. coli* strains. (**A-D**) Co-inoculation with Sym A and Δ*intS*. (**E-H**) Co-inoculation with Sym A and CmL05G13. (**I-L**) Co-inoculation with Sym A and Δ*cyaA*. (**A, E, I**) Proportion of adult insects with white symbiotic organ per rearing container, each containing 16-35 insects, indicating the level of *E. coli* predominance in the co-inoculated host insects. (**B, F, J**) Adult emergence rates of Sym A-inoculated, co-inoculated and *E. coli*-inoculated insects. (**C, G, K**) Bacterial titers in Sym A-inoculated, co-inoculated and *E. coli*-inoculated second instar nymphs in terms of bacterial *groEL* gene copies per nymph. (**D, H, L**) Bacterial titers in Sym A-inoculated, co-inoculated and *E. coli*-inoculated adult insects in terms of bacterial *groEL* gene copies per dissected symbiotic organ. In (**B-D**), (**F-H**) and (**J-L**), blue, gray, and orange indicate Sym A-inoculated (with adult emergence rates and bacterial titers shown in Fig. 2 and Fig. 3), co-inoculated, and *E. coli*-inoculated (with adult emergence rates and bacterial titers shown in Fig. 2 and Fig. 3), respectively. Bars, whiskers and dots indicate means, standard deviations and data points, respectively. Different alphabetical letters (a, b, c) indicate statistically significant differences (pairwise Wilcoxon rank-sum test with Hommel’s correction; *P* < 0.05).

#### Sym A vs evolved symbiotic *E. coli* CmL05G13

When Sym A cells and CmL05G13 cells were co-inoculated, CmL05G13 established predominant infection over Sym A at the second instar stage (Fig. 8A). The infection predominance of CmL05G13 continued until the adult stage, although Sym A infection recovered in a small fraction of the adult insects (Fig. 8B; Fig. 9E). The adult emergence rates of the co-inoculated insects were comparable to those of CmL05G13-infected insects and significantly lower than those of Sym A-infected insects (Fig. 9F), which probably reflects the preferential establishment of CmL05G13 in the co-inoculated adult insects (Fig. 8B). The bacterial titers in the co-inoculated insects were largely comparable to those in the CmL05G13-infected insects and the Sym A-infected insects (Fig. 9G, H). These results indicated that the evolved symbiotic *E. coli* CmL05G13 is superior to the cultivable symbiont Sym A in infection competitiveness, but inferior in host survival improvement. These features of CmL05G13 seem to be regarded as a “cheater” phenotype.

#### Sym A vs artificial symbiotic *E. coli* Δ*cyaA*

When Sym A cells and Δ*cyaA* cells were co-inoculated, Δ*cyaA* established predominant infection over Sym A at the second instar stage (Fig. 8A). However, the infection predominance was reversed at the adult stage, when Sym A became predominant over Δ*cyaA* (Fig. 8B; Fig. 9I). The adult emergence rates of the co-inoculated insects were comparable to those of the Sym A-infected insects and significantly higher than those of the Δ*cyaA*-infected insects (Fig. 9J), which probably reflects the predominant establishment of SymA in the co-inoculated adult insects (Fig. 8B). The bacterial titers in the co-inoculated insects were largely comparable to those in the Sym A-infected insects and the Δ*cyaA*-infected insects (Fig. 9K, L). These results indicated that, although the artificial symbiotic *E. coli* Δ*cyaA* may be good at initial infection and proliferation, the native uncultivable symbiont Sym A is superior in establishing the symbiosis with its original host *P. stali* and finally excludes the competitor *E. coli*, thereby attaining improved host survival.

## DISCUSSION

In this study, we experimentally investigated the symbiotic capabilities of the native uncultivable symbiont of *P. stali* (Sym A fixed in the Japan mainland populations) and the cultivable symbionts of *P. stali* (Sym C, Sym D, Sym E and Sym F present in the Ryukyu Islands populations) in comparison with the nonsymbiotic control *E. coli* (Δ*intS*), the evolved symbiotic *E. coli* (CmL05G13) and the artificial symbiotic *E. coli* (Δ*cyaA*) under the same host genetic background. Through single infection and competitive infection experiments, we attempted to address the intriguing question as to how *E. coli* strains that have evolved beneficial traits for *P. stali* in the laboratory perform symbiotically, either worse, equivalent, or better, in comparison with the native symbionts of *P. stali* that have evolved in nature.

In the single infection experiments, both the evolved and artificial symbiotic *E. coli* strains achieved adult emergence rates comparable to those of the cultivable symbionts, although they did not reach the level of the native uncultivable symbiont (Fig. 2A). These results demonstrate that the symbiotic *E. coli* strains can support host growth and survival at similar levels to the natural cultivable symbionts of *P. stali*. In contrast, both the evolved and artificial symbiotic *E. coli* strains exhibited evidently lower host fecundity than the cultivable symbionts of *P. stali* (Fig. 2B). These patterns are plausibly because of the previous experimental evolutionary setting that selected for improved adult emergence rate and body color (12), wherein fecundity was outside the scope of direct selection target. These results uncover that the symbiotic *E. coli* strains have evolved to contribute to different host fitness components in different ways.

In the competitive infection experiments, several remarkable properties of the symbiotic *E. coli* strains emerged. In summary, overall, the symbiotic *E. coli* strains usually, if not always, exhibited better competitive symbiotic capabilities than the natural symbionts of *P. stali*. Most notably, the evolved symbiotic *E. coli* strain CmL05G13 outcompeted all the cultivable symbionts Sym C, Sym D, Sym E and Sym F (Figs. 4-7), and furthermore, outcompeted the native uncultivable symbiont Sym A (Figs. 8-9). It was surprising that the native symbiont Sym A, which is genome-reduced, uncultivable, essential for host growth and survival, and fixed in natural host populations, was defeated and replaced by the co-infected symbiotic *E. coli* strain that evolved in the laboratory within a year or so. This finding reinforces the notion that the evolution of symbiosis can occur more easily and rapidly than conventionally envisaged (12, 22).

On the other hand, it should be noted that, despite the superior infection competitiveness, the symbiotic *E. coli* strains only poorly supported host’s fecundity and reproductive maturation in comparison with the natural symbionts of *P. stali* (Fig. 2B, C). Here, it is expected that, if the symbiotic *E. coli* were to infect a stinkbug already harboring a natural symbiont, it could potentially replace the essential symbiont and establish its own infection, while decreasing the host fitness. Theoretical and empirical studies have pointed out that mutualistic relationships are potentially prone to invasion by cheaters that obtain benefits from the relationship while avoiding associated costs (23–26). In this context, the symbiotic *E. coli* strain like CmL05G13 can be regarded as an “artificial cheater symbiont”, which may provide a unique experimental model to investigate the evolutionary dynamics of cheating and mutualism in symbiosis.

In this study, we found that the native *Pantoea* symbionts of *P. stali* are competitively replaced by the symbiotic *E. coli* strains, which are originally nonsymbiotic bacteria unrelated to *P. stali*, when co-inoculated. This striking finding looks contrasting to the previous reports on co-inoculation experiments using coreoid stinkbugs associated with *Caballeronia* symbionts, in which the native symbiont strains generally outcompete other potentially symbiotic bacterial strains and thereby establish the specific symbiotic associations (17, 18). Here, it should be noted that the symbiont transmission routes are different between the *P. stali*–*Pantoea* symbiosis and the coreoid–*Caballeronia* symbiosis. In *P. stali* and allied pentatomid stinkbugs, adult females vertically transmit their symbiotic bacteria to next generation nymphs by smearing *Pantoea*-containing excretion onto egg surface upon oviposition, for which female stinkbugs have evolved specialized histology, morphology and behavior (9, 10, 12, 27). By contrast, in *R. pedestris* and other coreoid stinkbugs, adult females usually do not transmit their symbiotic bacteria vertically, and newborn nymphs acquire the specific lineages of symbiotic *Caballeronia* from the environment every generation (28–30). Since the environmental symbiont acquisition observed in *R. pedestris* and other coreoid stinkbugs inevitably entails nymphal oral acquisition of diverse microbial species, it seems likely that natural selection has favored the evolution of such mechanisms that the native beneficial symbiont strain can specifically colonize and establish infection in the host symbiotic organ (31, 32). On the other hand, in the vertical symbiont transmission system of *P. stali*, since the native symbiotic bacteria are purely smeared on eggshell and ready for oral acquisition by newborn nymphs, the natural selection that favors the mechanisms for ensuring selective colonization and establishment of the native symbiotic bacteria may be relaxed. In this context, although speculative, we suggest the possibility that, in the native symbiotic bacteria of *P. stali* that have experienced continuous vertical transmission and long-lasting coevolution with their host insect, fitness contribution to the host insect rather than infection competitiveness has been optimized, which may constitute the basis for the observation that the native symbiotic bacteria were competitively defeated by the symbiotic *E. coli* strains.

It remains an intriguing yet unresolved question what mechanisms enable the evolved symbiotic *E. coli* to enhance the host’s emergence rate and to outcompete the native symbiotic bacteria. Considering that symbiont titers tend to show positive correlations to host fitness levels among the uncultivable and cultivable symbiotic bacteria as well as among the nonsymbiotic and symbiotic *E. coli* strains (Figs. 2 and 3), quantitative effects via increased bacterial titers appear plausible. As for qualitative effects, comparative transcriptomic, proteomic, and metabolomic analyses of the experimental *E. coli* strains as well as the host symbiotic organ harboring them before and after the symbiotic evolution may provide important clues. It is notable that, while the evolved symbiotic *E. coli* strain CmL05G13, which accumulated numerous mutations including Δ*cyaA* during the experimental evolution (12), outcompeted the native symbiont Sym A, the artificially engineered symbiotic *E. coli* strain Δ*cyaA*, in which only *cyaA* gene was disrupted (12), failed to replace SymA (Figs. 8, 9). These results strongly suggest that additional mutations other than Δ*cyaA* accumulated in the evolved symbiotic *E. coli* strain CmL05G13 also contribute to the enhanced symbiotic performance. Identification of these mutations and characterization of their roles in symbiosis represent an important avenue for future research.

Here we note that our experimental studies are performed in the laboratory under artificial settings, and thus these results should be carefully treated in making biological, ecological and evolutionary inferences. Co-inoculation experiments were conducted using drinking water suspending 1:1 bacterial mixture at a standardized cell density, but, considering the relative dominance of the native symbiont on dry eggshell compared to contaminated bacterial competitors, the competitive “start lines” must be quite different in the field. For standardized comparison, we used a single insect genotype and a limited number of bacterial genotypes, but both the host insects and the bacterial associates are genetically diverse in the real world. We evaluated fitness contribution of the bacterial infections in terms of adult emergence rate and 30-day fecundity, but such important measures as lifetime reproduction, offspring quality and transgenerational effects remain to be quantified.

In conclusion, we demonstrate that the artificially evolved symbiotic *E. coli* strains obtained in the laboratory can surpass the natural symbiotic bacteria of the stinkbug *P. stali* in terms of competitive colonization ability. It is striking that *E. coli*, originally a gut bacterium of vertebrates with no natural association with the insect, can rapidly evolve such a remarkable symbiotic capability. On the other hand, the fitness effects of the symbiotic *E. coli* strains on host’s reproductive success are inferior in comparison with the natural symbiotic bacteria of *P. stali*, which constitute a “cheater-like” property of the symbiotic *E. coli* strains. Taken together, these results highlight that the artificially constructed *P. stali*–*E. coli* experimental symbiotic system serves as a powerful model for empirical understanding of the mechanisms and dynamics of the evolution of symbiosis.

## MATERIALS AND METHODS

### Host insect strain

A long-term laboratory strain of the brown-winged green stinkbug *P. stali*, originally derived from adult insects collected in Tsukuba, Ibaraki, Japan in September 2012, was used. Early-instar nymphs were reared in sterilized disposable plastic dishes (90 mm in diameter, 20 mm in height) lined with filter paper and supplied with two raw peanuts and cotton balls soaked in water supplemented with 0.05% (w/v) ascorbic acid (DWA). From the third instar onward, nymphs were transferred to larger plastic containers (150 mm in diameter, 60 mm in height) and provided with a sufficient quantity of raw peanuts and DWA. All insects were maintained under a long-day photoperiod (16 h light: 8 h dark) at 25 ± 1 °C and 50 ± 5% relative humidity (20, 33).

### Symbiont strains

The uncultivable symbiont Sym A was freshly prepared as needed by dissecting and homogenizing the laboratory-maintained insects of *P. stali*. The cultivable symbionts Sym C, Sym D, Sym E and Sym F were stock cultures that had isolated from field populations of *P. stali* as previously described (9).

### E. coli strains

The following *E. coli* strains were used in this study. Δ*intS* is an ordinary *E. coli* strain with little symbiotic capability, which was used as a nonsymbiotic control *E. coli* strain. CmL05G13 is an evolved symbiotic *E. coli* strain obtained through a previous experimental symbiotic evolution, in which a hypermutating *E. coli* strain Δ*mutS* was continuously selected for improved adult body color for 13 host generations (12). The symbiotic CmL05G13 strain accumulated many mutations during the experimental symbiotic evolution, of which a disruptive mutation in *cyaA* gene encoding adenylate cyclase was identified as a major effect gene responsible for its mutualistic phenotypes (12). In this context, Δ*cyaA* was used as an artificial symbiotic *E. coli* strain, which was obtained from the Keio *E. coli* single gene knockout collection (34).

### Preparation of aseptic insects

To obtain symbiont-free aseptic nymphs of *P. stali*, egg masses were collected and surface-sterilized by immersing in 4% formaldehyde for 10 min twice, followed by rinsing with sterilized water for 10 min twice, and then air-dried. The sterilized egg masses were stored in groups of three per plastic Petri dish at 25 ± 1 °C until use. Raw peanuts were immersed in 99% ethanol for 30 min, vacuum-dried, and stored under refrigeration. Filter papers were autoclaved at 121°C for 20 min and dried in a drying oven. These sterilized materials were used for maintaining aseptic nymphs and for performing bacterial inoculation as described previously (20).

### Bacterial inoculation to aseptic insects

*E. coli* strains Δ*intS*, CmL05G13 and Δ*cyaA*, as well as the cultivable symbiont strains Sym C, Sym D, Sym E and Sym F, were each cultured overnight at 25°C with shaking in 3 mL of Luria-Bertani (LB) liquid medium (1% tryptone, 0.5% yeast extract, 1% NaCl). Subsequently, 200 µL of each overnight culture was transferred into 2.8 mL of fresh LB medium and cultured with shaking for 5 h. The cultures were then diluted with sterilized water to optical density at 600 nm (OD_600_) to be 0.1, which were used as bacterial inocula.

Within 24 h after hatching from each surface-sterilized egg mass, newborn nymphs were kept in a plastic Petri dish without water for a day. Thereafter, two cotton balls, each soaked with 0.75 mL of the inoculum, were placed in each dish to allow the nymphs to ingest the bacterial suspension. Note that OD_600_ = 0.1 corresponds to approximately 1 × 10^5^ colony formation units (CFUs) per mL for the *E. coli* strains as well as the cultivable symbiont strains, and therefore the inoculum is estimated to contain 1 × 10^5^ bacterial cells per mL.

The uncultivable symbiont Sym A was prepared by dissecting the symbiotic organs from three female adults under a stereomicroscope (Leica, S9i) in phosphate buffered saline (PBS: 137 mM NaCl, 8.1 mM Na_2_HPO_4_, 2.7 mM KCl, 1.5 mM KH_2_PO_4_, pH 7.4) using tweezers and scissors. Each sample was homogenized in 200 µL of PBS, and 60 µL of the homogenate was diluted with 1,440 µL of sterilized water to prepare the inoculum for one Petri dish. The number of SymA cells was estimated by counting bacterial cells prepared from five symbiotic organs of female adults using a hemocytometer, which yielded the value of approximately 5 × 10^8^ bacterial cells per organ. Therefore, the inoculum contained approximately 1 × 10^5^ cells/mL.

Competitive infection assays were performed by mixing two inocula (750 µL each) at a 1:1 ratio and inoculating the mixture to aseptic nymphs orally as described above. The bacterial combinations tested were as follows: (1) the cultivable symbiont Sym C versus the control nonsymbiotic *E. coli* Δ*intS*, the evolved symbiotic *E. coli* CmL05G13, or the artificial symbiotic *E. coli* Δ*cyaA*; (2) the cultivable symbiont Sym D, Sym E, or Sym F versus the evolved symbiotic *E. coli* CmL05G13; and (3) the uncultivable SymA versus the control nonsymbiotic *E. coli* Δ*intS*, the evolved symbiotic *E. coli* CmL05G13, or the artificially symbiotic *E. coli* Δ*cyaA*.

### Bacterial quantification

Bacterial quantification was performed at two developmental stages, namely the third day of the second instar nymphal stage and the adult stage. Quantification of the cultivable bacterial strains was carried out by CFU counting. On the third day of the second instar nymphal stage, each insect was homogenized in 200 µL PBS, the homogenate was diluted 10^-2^, 10^-3^ and 10^-4^ times with sterilized water, and 100 µL of aliquot for each dilution was plated onto a LB agar plate. Following an overnight incubation at room temperature, colonies were counted. For adults, each symbiotic organ dissected from an adult insect was homogenized in 200 µL of PBS, diluted 10^-4^, 10^-5^ and 10^-6^ times with sterilized water, and plated in the same manner for colony counting.

Quantification of the uncultivable symbiont Sym A and the competitor *E. coli* strains was performed by quantitative PCR. Each whole second instar nymph or dissected adult symbiotic organ was homogenized in a plastic tube with a pestle in 200 µL of PBS and subjected to alkaline DNA extraction essentially as described previously (35). A 10 µL aliquot of the homogenate was transferred to a new tube and mixed with 32 µL of alkaline solution (25 mM NaOH, 0.2 mM EDTA), followed by heating at 95°C for 10 min. The solution was then neutralized by adding 64 µL of neutralizing solution (12.5 mM HCl, 10 mM Tris) and placed on ice. The DNA samples were centrifuged and subjected to quantitative PCR either directly or after a 100-fold dilution with sterilized water. For Sym A, *groEL* gene was targeted using the primers AgroL1013F (5′-ATC AGG GCG CAA TAT CTG GT-3′) and AgroL1103R (5′-CGC TCC TGC AGT TTC TCT TTG-3′), which amplify a 91 bp fragment. For *E. coli*, *groEL* gene was also targeted using the primers EgroL344F (5’-ACC TGA AAC GTG GTA TCG AC-3’) and EgroL450R (5’-TTT CGT CGG AGT TAG CGG AG-3’), which amplify a 126 bp fragment. The quantitative PCR reaction mixtures consist of 300 nM each of forward and reverse primers, 2 × KAPA SYBR FAST qPCR Master Mix Universal (KAPA Biosystems), 5 μl of the sample solution, and sterilized water, yielding a total volume of 20 μl. Quantitative PCR was performed using a Stratagene Mx3000P (Agilent Technologies) under the following thermal profile: 95°C for 3 min followed by 40 cycles of 95°C for 5 sec and 58°C for 15 sec. Each sample was quantified in duplicate. Standard curves were constructed using a linearized plasmid containing the target site, with concentrations corresponding to 5.45 × 10^2^, 10^3^, 10^4^, 10^5^, 10^6^, and 10^7^ *groEL* gene copies for Sym A, and 6.3 × 10^2^, 10^3^, 10^4^, 10^5^, 10^6^, and 10^7^ *groEL* gene copies for *E. coli*.

## ACKNOWLEDGMENTS

We thank Tomoko Matsushita for support of rearing and supply of experimental insects.

## FUNDING

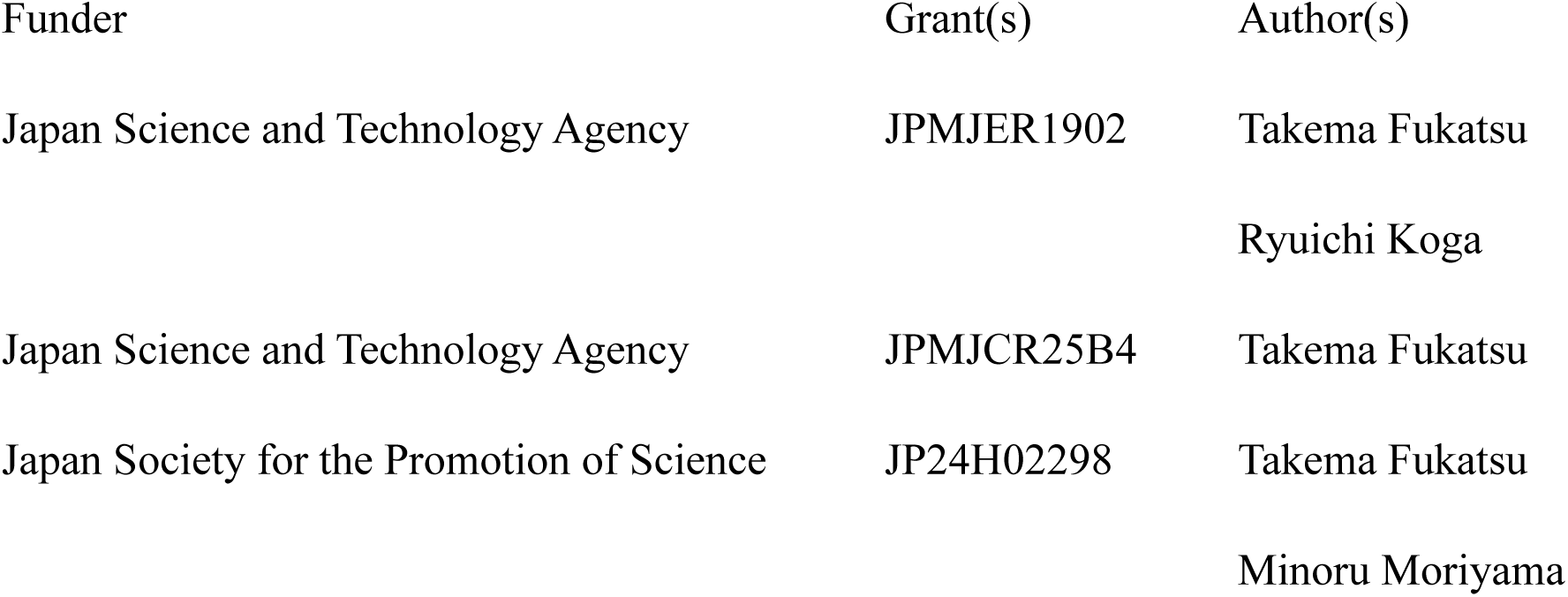

## AUTHOR CONTRIBUTIONS

Wenjin Cai, Conceptualization, Formal analysis, Investigation, Writing – original draft | Minoru Moriyama, Formal analysis, Investigation, Writing – review and editing | Yudai Nishide, Investigation, Writing – review and editing | Ryuichi Koga, Project administration, Writing ‒ review and editing | Takema Fukatsu, Conceptualization, Funding acquisition, Project administration, Supervision, Writing – original draft, Writing ‒ review and editing

## DIRECT CONTRIBUTION

This article is a direct contribution from Takema Fukatsu, a Fellow of the American Academy of Microbiology, who arranged for and secured reviews by Kazutaka Takeshita, Akita Prefectural University, and Hiroaki Noda, National Agriculture and Food Research Organization (NARO).

## DATA AVAILABILITY

All the source data for constructing Figs. 2–9 are provided in Supplemental Table S1.

## ADDITIONAL FILES

Supplemental Table S1.

